# Strainline: full-length de novo viral haplotype reconstruction from noisy long reads

**DOI:** 10.1101/2021.07.02.450893

**Authors:** Xiao Luo, Xiongbin Kang, Alexander Schönhuth

## Abstract

Haplotype-resolved de novo assembly of highly diverse virus genomes is critical in prevention, control and treatment of viral diseases. Current methods either can handle only relatively accurate short read data, or collapse haplotype-specific variations into consensus sequence. Here, we present Strainline, a novel approach to assemble viral haplotypes from noisy long reads without a reference genome. As a crucial consequence, Strainline is the first approach to provide strain-resolved, full-length de novo assemblies of viral quasispecies from noisy third-generation sequencing data. Benchmarking experiments on both simulated and real datasets of varying complexity and diversity confirm this novelty, by demonstrating the superiority of Strainline in terms of relevant criteria in comparison with the state of the art.

## Background

Viruses such as HIV, ZIKV and Ebola lack proofreading mechanisms when they replicate themselves with RNA-dependent RNA polymerase (RdRp) Holland et al. (1992); Domingo et al. (1996). Therefore, they are characterized by high mutation rates, and commonly populate hosts as a collection of closely related strains which differ by only small amounts of variants, and which together are referred to as viral quasispecies Domingo et al. (2012). The genetic diversity of viral quasispecies plays an important role in viral evolution. Among others, it contributes to tissue tropism, virus transmission, disease progression, virulence and drug/vaccine resistance *Holland et al.* (1992); Beerenwinkel et al. (2005); Douek et al. (2006); Knyazev et al. (2021). In addition, biological functionalities or phenotypic appearance can differ substantially across different strains *Loman et al.* (2013). Currently, the Covid-19 pandemic puts the necessity to monitor the outbreak of viruses, to track their evolutionary history, and to develop effective vaccines and drugs in the spotlight of greater public interest. To accurately account for these issues, accurate reconstruction of strain-resolved genomes is a prerequisite.

It is the general, ultimate goal of viral quasispecies assembly to reconstruct the individual, strain-specific haplotypes at their *full length*. Further, along with strain identity-preserving sequence, accurate *estimates of strain abundances* are required for full quantification of infections at the RNA/DNA level. Notwithstanding the short size of virus genomes, it is still a challenge because within a viral quasispecies (i) closely related strains share plenty of near-identical genomic fragments, single strains are affected by repetitive regions Somerville et al. (2019), (iii) the number of strains is unknown, and (iv) the abundances of strains vary across the strains, which is further aggravated by read coverage fluctuations along the genomes.

So far, existing methods for viral quasispecies assembly can be classified into *reference-based* approaches on the one hand and *de novo* (reference free) approaches on the other hand. Reference-based methods such as ShoRAH Zagordi et al. (2011) and PredictHaplo Prabhakaran et al. (2013) require high quality reference for reliable reconstruction of strains and have been specializing in processing relatively error-free short read data. Importantly, high quality reference genomes may not be available precisely when they are needed the most: very often, new outbreaks of known viruses are caused by virus variants that significantly deviate from curated reference sequence. Last but not least, reference-guided methods are prone to introducing biases and can be blind with respect to crucial variant-related details in genomic regions of particular interest Töpfer et al. (2014); Baaijens et al. (2017).

*De novo* (reference free) viral quasispecies assembly tools, such as SAVAGE Baaijens et al. (2017) or viaDBG Freire et al. (2021), both are able to employ overlap and de Bruijn graph-based techniques to assemble NGS reads into haplotype-specific contigs (a.k.a. haplotigs), where the two assembly paradigms, overlap vs. de Bruijn graph based, come with different advantages and disadvantages. The resulting contigs of these short read based approaches tend to be too short to span genomes at their full length. The reason are sequence patches that are shared by different strains (and also repetitive areas within strains), which induce ambiguities that cannot be overcome by short reads themselves. For computing full-length genomes, one can try to leverage the strain-specific abundances, which allows to bridge contigs across otherwise ambiguous stretches of sequences. To this end, methods such as Virus-VG Baaijens et al. (2019) and VG-Flow Baaijens et al. (2020) have been developed, the latter approach of which introduced flow variation graphs as a computational concept of potential greater value. Because the runtime is polynomial in the length of the genomes, VG-Flow Baaijens et al. (2020) can also be used for bacteria sized genomes.

We recall that all these existing approaches focus on viral haplotype reconstruction from *short and accurate next-generation sequencing (NGS) reads*, as generated most prominently by Illumina platforms. Again, the fact that short reads fail to span inter- and intra-genomic identical regions crucially hampers the process of reconstructing full-length viral haplotypes. Leveraging strain-specific abundances, as implemented by VG-Flow Baaijens et al. (2019) for example, are not necessarily able to output full-length, strain-specific assembled sequence for certain viruses, such as ZIKV and Polio.

*Quite apparently, virus genome assembly methods have approached their limits when operating with short read NGS data.* Processing long and noisy third-generation sequencing (TGS) data, such as generated by Pacific BioSciences (PacBio), performing single-molecule real-time (SMRT) sequencing, and Oxford Nanopore (ONT), performing nanopore sequencing, as the currently two most popular sequencing platforms, offer rescue.

The length of TGS reads ranges from several Kbp to hundreds of Kbp, or even to ~Mbp Logsdon et al. (2020). TGS reads enable to span intra-genomic repeats and areas shared by different genomes, hence cover regions that are unique to single strains Somerville et al. (2019). So, in comparison with NGS reads, TGS reads have considerably greater potential to resolve ambiguities across different strains. The drawback of TGS reads are the elevated error rates they are affected with. Unlike for NGS platforms (sequencing error rate < 1%), error rates of PacBio CLR and ONT reads range from 5 to 15%, which raises the issue of sequencing errors to a greater order of magnitude.

There are a handful of *de novo* assembly methods that specialize in processing error-prone long reads such as FALCON Chin et al. (2016), Canu Koren et al. (2017), Flye Kolmogorov et al. (2019), Wtdbg2 Ruan and Li (2020) and Shasta Shafin et al. (2020), all of which have been published fairly recently. None of these approaches makes a decided attempt to generate haplotype-(strain-) resolved genomic sequence. Rather, these approaches choose to output *consensus* sequence, as a summary across several or all haplotypes/strains in the mix. In other words, all of the assemblers presented in the literature so far fall under the category ‘generic (or consensus) assembler’.

In addition, metaFlye, originally designed to perform assembly of metagenomes, operates at the level of species Vicedomini et al. (2021), so neglects to resolve individual genomes at the level of strains.

In conclusion, haplotype-aware assembly of viral quasispecies from erroneous long reads can still be considered an issue that has remained unresolved so far: no method is able to address the issue satisfyingly.

Here, we pursue a novel strategy to resolve the issue. In that, our approach is the first one to accurately reconstruct the haplotypes of viral quasispecies from third-generation sequencing reads, to the best of our knowledge. We recall that processing long TGS reads appears to be the only current option to reconstruct genomes at the level of strains, for the majority of the currently predominant viruses.

In a brief description (see below for details), our novel strategy consists of *local de Bruijn graph based assembly* in a first step that addresses to wipe out errors. Subsequently, we turn our attention to an overlap graph-based scheme by which to iteratively extend haplotype-specific contigs (haplotigs) into full-length haplotypes. After a filtering step that removes artifacts and preserves true sequence, our approach outputs a set of haplotypes—a large fraction of which appear to have reached full length—together with the relative abundances of the haplotypes within the mix of haplotypes.

We evaluate our approach on various virus datasets that have been approved earlier in the literature. For each dataset, we process both PacBio CLR reads and ONT reads, as the two most predominant types of TGS reads. Benchmarking results on both simulated and real data confirm our claims: our approach accurately reconstructs all full-length haplotypes and delivers sufficiently accurate estimates of their relative abundances.

We also compare our approach with the current state of the art. We recall however that none of the current approaches decidedly addresses the issue of strain-resolved viral quasispecies assembly from long reads. As a consequence, our approach outperforms the state of the art rather drastically. Our approach has its greatest advantages in terms of haplotype coverage, reaching nearly 100% on all datasets, where other methods never get beyond 60-70% (if they get there at all; in particular on ONT data, alternative approaches the limit is reached substantially earlier for other approaches). Further marked advantages are assembly contiguity (measured as per N50 or NGA50) and accuracy (expressed by low error rates and little misassembled contigs). A currently particularly interesting application scenario is the assembly of haplotype-resolved genomes of SARS-CoV-2, because strain resolution will sharpen our understanding about mutation rates and evolutionary development of the virus. Also in this scenario of particular current interest, our approach demonstrates to outperform all existing approaches by fairly large margins.

## Results

We have designed and implemented Strainline, a novel approach that implements the strategy that we already sketched above, and will be described in full detail in the following. In a short summary, Strainline reconstructs full-length, strain-resolved viral haplotypes from noisy long read (TGS read) sequencing data. Strainline is a *de novo* assembler, so does not have to rely on available reference sequence. Therefore, Strainline operates free of biases induced by prior knowledge, which has been repeatedly pointed out as a source of issues in earlier work.

In this section, we provide a high-level description of the workflow and evaluate its performance on both simulated and real data, in comparison with existing state-of-the-art tools. In our comparisons, we focus on both generic and metagenome assembly approaches that are able to process long reads without having to rely on reference sequence, which matches the conditions under which Strainline is able to operate.

### Approach

See Figure 1 for an illustration of the overall workflow of Strainline. Here, we describe the workflow briefly. For detailed descriptions of the individual steps, we refer to the Methods section.

**Figure 1.**
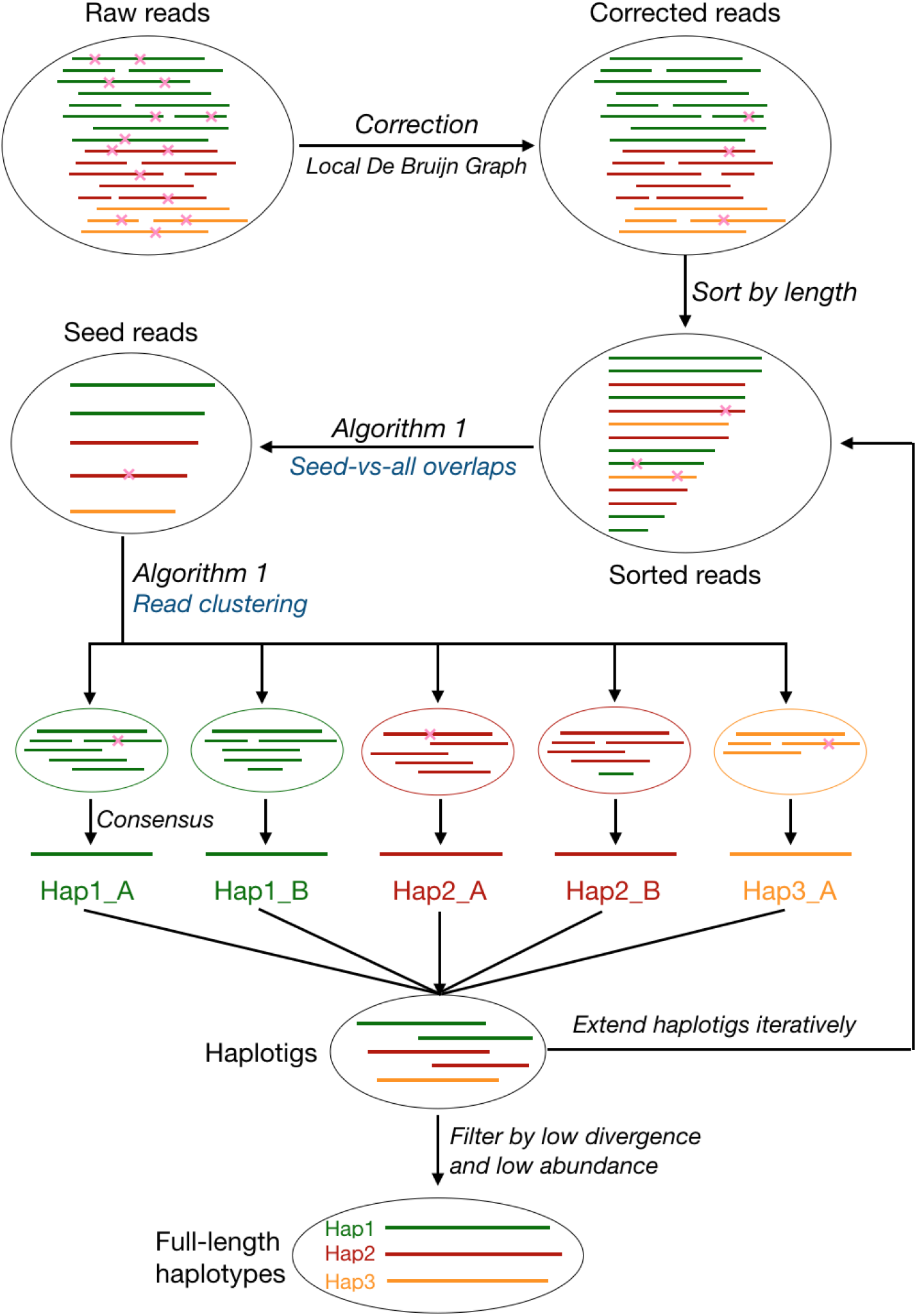
The workflow of Strainline. The reads with different colors are from different haplotypes or strains. The pink fork represents the sequencing error, i.e. mismatch, insertion or deletion. The two steps *Seed-vs-all overlaps* and *Read clustering* are executed simultaneously (see Algorithm 1 for details). Hap1 A and Hap1 B denote two subsequences (not full-length) of haplotype 1 (Hap1), the same for Hap2 A and Hap2 B. There still may be very few remaining sequencing errors in corrected reads such as in the corresponding read cluster of Hap1 A, and incorrectly clustered reads such as the ‘green read’ in the corresponding read cluster of Hap2 B. Nevertheless, these errors will be eliminated through *Consensus* step.

Strainline consists of three stages. The first stage addresses to correct sequencing errors in the raw long reads, for which it employs *local De Bruijn graph assembly*. The second stage addresses to iteratively extend haplotype-specific contigs into full-length haplotype-specific genomes, based on an overlap based strategy. The third stage, finally, is for filtering the resulting contigs so as to remove haplotypes of too low divergence in comparison with others (so likely reflect errors instead of strain-specific variation), or too low abundance (so likely reflect artifacts). The eventual output is a set of full-length haplotypes along with their corresponding relative frequencies, clear of errors and artifacts.

In general, De Bruijn graph based approaches tend to be inappropriate for TGS read data, because of the elevated error rates that apply. However, we found a (local) de Bruijn graph based approach that was originally developed for long genomes, to effectively work for genomes of sizes of that corresponded to viruses (that is in the tens of thousands of nucleotides) Tischler and Myers (2017). Apparently, the superiority of the approach when dealing with virus sized genomes had passed unnoticed earlier.

Given a target read to be corrected, the corresponding strategy considers the reads that overlap the target read, where overlaps are determined based on evaluating canonical k-mers. The resulting overlapping reads together with its target read form a read alignment pile that is divided into small windows. Subsequently, a De Bruijn graph is constructed for each such small window. Based on evaluating the graph, an optimal consensus sequence is determined, which reflects the error corrected, true sequence of the target read. For details, see Figure 2.

**Figure 2.**
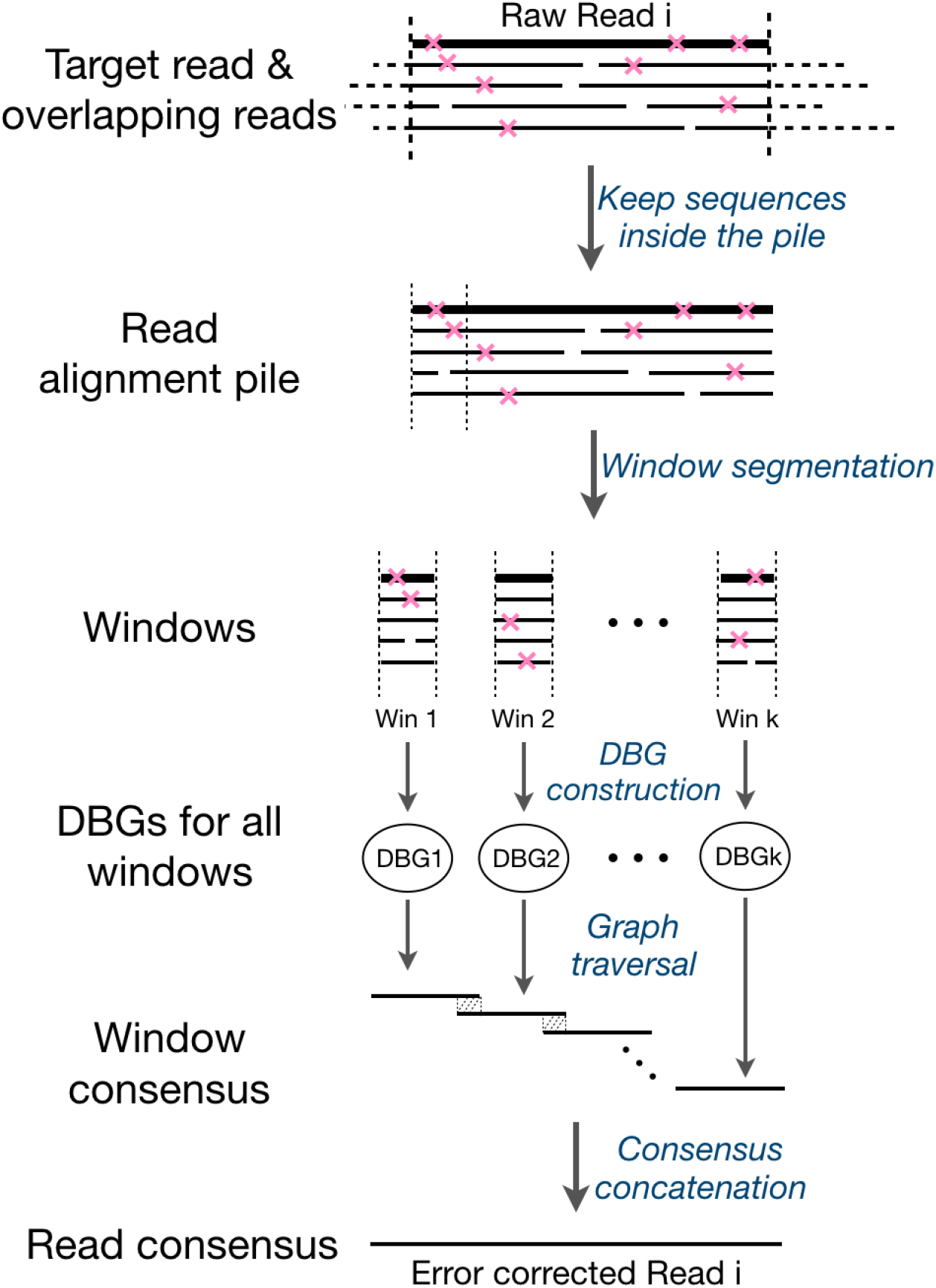
The schematic diagram for the sequencing error correction procedure of raw reads. In the top region, the bold solid line denotes the target raw read *i*, and the overlapping reads of the target read *i* are drawn dashed outside of the read alignment pile and solid inside of it. The pink fork represents the sequencing error, i.e. mismatch, insertion or deletion. The read alignment pile is split into *k* small windows, representing as Win 1, Win 2, … Win *k*. DBG is short for De Bruijn Graph. The region filled with oblique dashed lines between two window consensus denotes the overlap between them.

The second stage falls into two sub-steps. Firstly, Strainline determines read clusters where each of the clusters reflects a collection of reads that overlap each other in terms of genomic position (computed by ‘Algorithm 1’ in Figure 1). Secondly, Strainline uses a partial order alignment (POA) algorithm to yield a consensus sequence for each read cluster. These consensus sequences are expected to be haplotype-specific contigs (haplotigs). The haplotigs are then iteratively extended into full-length haplotypes.

The third and final stage is to filter haplotypes having very low divergence or very low relative abundance. Most likely, such haplotypes were introduced due to redundant or spurious sequences. See Methods for full details on each of the stages involved in the overall workflow.

### Datasets

#### Simulated data

We simulated various datasets using both PacBio and ONT long read sequencing technologies, yielding the common (PacBio CLR and ONT), high sequencing error rates (5% ~15%), using PBSIM Ono et al. (2013) and NanoSim V2.6.0 Yang et al. (2017), as approved read simulators. We used four virus mixture datasets (HIV, Poliovirus, HCV and ZIKV), similar in terms of composition of strains to those presented by Baaijens et al. (2019). We further generated one additional dataset reflecting a SARS-CoV-2 quasispecies, composed of 5 strains (note that the number of SARS-CoV-2 strains that affect one individual is entirely unclear at this point, because of the lack of analysis tools). Notably, it is very common to perform ultra-deep sequencing supported by the short length of the viral genome (7~30kbp in our cases) Baaijens et al. (2019, 2017); Giallonardo et al. (2014). Therefore, we uniformly set the overall sequencing depth (the sum of average depth of each strain) as 20000 on all simulated datasets.

Additionally, in order to evaluate the effect of sequencing coverage, we generated 5-strain HIV datasets at an error rate of 10% and varying coverage, of overall depths of 500×, 1000×, 2000×, 5000×, 10000× and 20000×, respectively, while relative frequencies of strains did not change. More details about simulated data are shown in Data simulation.

##### 5-strain HIV mixture

This dataset consists of five known HIV-1 strains (YU2, NL43, JRCSF, HXB2, 896), as originally presented in Giallonardo et al. (2014). Strains were simulated at relative abundances between 10% and 30%, i.e. a sequencing coverage of 2000 to 6000 per strain. The pairwise divergence (here defined as 1 BLAST identity, where BLAST identity is the number of matching bases divided by the length of the BLAST alignment) varies from 2.6% to 6.9% between strains. This virus mixture is one of the most challenging datasets, because the highly repetitive regions in the HIV genome, which usually hamper the performance of short read based assemblers Giallonardo et al. (2014); Baaijens et al. (2017, 2019).

##### 6-strain Poliovirus mixture

This mixture contains six strains of Poliovirus (Type 2), with exponentially increasing relative abundances from 2% to 50%. The haplotype sequences were downloaded from the NCBI database. The pairwise divergence varies from 1.2% to 7.2%.

##### 10-strain HCV mixture

This mixture contains ten strains of hepatitis C virus (HCV), Subtype 1a, with relative frequencies varying from 5% to 15% per haplotype. The haplotype sequences were also obtained from the NCBI database, with a pairwise divergence of 2.7% to 7.5%.

##### 15-strain ZIKV mixture

This mixture consists of fifteen strains of Zika Virus (ZIKV), of which three master strains were obtained from the NCBI database and four mutants were generated per master strain by randomly introducing mutations. The pairwise divergence changes from 1.0% to 15.8% and the relative frequency varies between 2% and 12%.

##### 5-strain SARS-CoV-2 mixture

This mixture consists of five strains of SARS-CoV-2, with the relative frequencies varying from 10% to 30%. The true haplotype sequences (high quality without N bases) were extracted from different regions (namely, Belgium, Egypt, Oman, USA, China) in the GISAID (https://www.gisaid.org/) database, with a pairwise divergence of 2.2% to 14.5%.

#### Real data

To evaluate our method on real sequencing data, we downloaded two real datasets for benchmarking analysis.

##### 5-strain PVY mixture

This dataset is consist of five strains Potato virus Y (PVY). The true sequences of five strains were accessed from GenBank under accession numbers MT264731–MT264741. The pairwise divergence varies from 3.5% to 21%. The corresponding real ONT reads were obtained from the SRA database under BioProject PRJNA612026, as recently presented in Della Bartola et al. (2020). We downloaded long read sequencing data for each strain and then mixed them together to generate a pseudo virus mixture, where strains have relative frequencies varying from 9% to 32% and the total sequencing depth is approximate 5800×.

##### SARS-CoV-2 real sample

This dataset is Oxford Nanopore sequencing (GridION) data of a real SARS-CoV-2 sample which is downloaded from SRA database: SRP250446. The N50 of the length of the reads is 2.5kbp, the average sequencing error rate is approximately 10% and the average sequencing coverage about 12000×.

### Benchmarking: Alternative Approaches

We recall that Strainline is unique insofar as it is the first approach to determine the haplotype/strain-specific genomes of viruses from long reads *de novo.* For the sake of a meaningful comparison, we chose long read de novo assemblers that are designed to deal with mixed samples (in other words, designed for metagenome assembly), such as Canu Koren et al. (2017) and metaFlye Vicedomini et al. (2021), on the one hand, and generic (consensus) de novo assemblers, such as wtdbg2 Ruan and Li (2020) and Shasta Shafin et al. (2020) on the other hand. Of those, we subsequently excluded metaFlye, because it failed to perform the assemblies. failed to run metaFlye on our datasets ^1^. Shasta returned too many fragmented contigs, indicating that no real assembly was computed. For fairness reasons—we recall that all tools were originally designed for different purposes—we excluded metaFlye and Shasta from further consideration. For Canu, we used the parameters recommended for metagenome assembly, and we ran wtdbg2 with default parameters. The output contigs were then subject to being evaluated.

### Performance Evaluation

#### Assembly Metrics

In the evaluation, we considered all relevant categories, as output by QUAST V5.1.0 Mikheenko et al. (2018), as a prominent assembly evaluation tool. As is common, we discarded contigs of length less than 500bp from the output of all tools. In particular, we ran the `metaquast.py` program with the option `--unique-mapping` appropriately taking into account that our data sets reflect mixed samples. In the following, we briefly define the metrics we are considering.

##### Haplotype coverage (HC)

Haplotype coverage is the percentage of aligned bases in the ground truth haplotypes covered by haplotigs. Haplotype coverage is commonly used to measure the completeness in terms of genome diversity of the assembly.

##### N50 and NGA50

We also consider N50 and NGA50 to measure assembly contiguity, as per their standard definitions: N50 is the maximum value such that all contigs of at least that length cover at least half of the assembly, and NGA50 is the maximum value such that the contigs of at least that length cover at least half of the reference sequence when aligned against it (after breaking contigs at misassembly events and trimming all unaligned nucleotides); here, the reference sequence is taken to reflect the concatenation of all strain specific genomes from which reads were simulated, or the canonical choices of reference sequences for real data otherwise.

##### Error rate (ER)

The error rate is the fraction of mismatches, indels and N’s (i.e. ambiguous bases) in the alignment of the contigs with the reference sequences.

##### Misassembled contigs proportion (MC)

If a contig involves at least one misassembly event, it is counted as *misassembled contig*. A misassembly event is given when contigs align with a gap or overlap of more than 1kbp, align to different strands, or even different haplotypes. As MC, we report the percentage of misassembled contigs relative to the overall number of output contigs.

##### Precision and recall

Because it was found helpful in evaluating virus genome assemblies earlier, we also report precision and recall. While precision refers to the fraction of contigs that align to the correct strain-specific sequence, recall refers to the fraction of strains that have a correctly aligned contig. Therefore, the edit distance of the alignment of the contig with the reference sequence must not exceed a threshold *d*, which can vary (e.g. 0%, 1%, 2%, 3%, 4%, 5%, see Figures S1-S3). In Results in the main text, the most stringent threshold 1% is used).

#### Haplotype abundance evaluation

Furthermore, we use two further metrics equally suggested in prior work as helpful for evaluating virus genome assembly quality Baaijens et al. (2019, 2020). Namely, we report the absolute frequency error (AFE) and the relative frequency error (RFE), which measure the deviation of the estimated abundances from the true abundances of the haplotypes. Let *k* be the number of true haplotypes. For a haplotype *i* ∈ [*k*], let 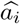 and *a_i_* be the estimated and true abundance of haplotype *i*, respectively. To determine 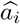, we first collect all haplotigs that get aligned with *i*, and then add up the abundances of these haplotigs. Let further 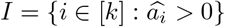 be all haplotypes that by their abundance were estimated to exist. One then calculates: 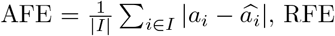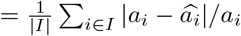

### Benchmarking results

We performed benchmarking experiments including all methods on the simulated and real data as described above, for both PacBio CLR and Oxford Nanopore reads. In a short summary of the results *ex ante*, Strainline manages to reconstruct all full-length haplotypes accurately from the mixed viral samples. Because truly specializing competing approaches are lacking, Strainline performs best in comparison with any alternative approach, by quite drastic margins.

#### Simulated PacBio CLR data

See Table 1. Strainline yields near-perfect assemblies on all five datasets (HIV, Poliovirus, HCV, ZIKV and SARS-CoV-2). That is, all ground truth haplotypes in the mixed samples are reconstructed to their full extent (HC≈100%, recall=100%), at very low error rates (0.002% ~ 0.074%) and without misassemblies. It also obtains the exact number of true haplotypes on four datasets (precision is 100%), and overestimates the number of haplotypes in the 5-strain SARS-CoV-2 mixture (precision is 71.4%). Note that the SARS-CoV-2 genome is particularly challenging because of its great length (30kbp) and the content of regions shared by different strains. In comparison, both Canu and Wtdbg2 struggle to reconstruct the haplotypes on all five datasets, with Canu achieving only 55.7% to 85.9% haplotype coverage on these datasets. Canu further achieves 100% recall on SARS-CoV-2, but only 60% to 80% recall on the other four datasets (see Figure S1). Notably, Strainline outperforms other tools in terms of precision on four datasets except that Canu achieves better precision on the 5-strain SARS-CoV-2 dataset. The reason for Canu’s successes in terms of precision and recall is the fact that its assembly is heavily fragmented (Canu generates 16 fragmented contigs, at an N50 of 12419), which puts the successes into a different context (and indicates that precision and recall have to be taken with a certain amount of care in the evaluation of assembly performance). Wtdbg2 basically generates a single consensus genome sequence rather than keeping haplotype information. Consequently, it merely obtains 10.7% to 20.6% haplotype coverage at high error rates, for example, 1.8%, 5.1%, 1.7% on HIV, HCV, ZIKV datasets, respectively.

**Table 1.** Benchmarking results for simulated PacBio CLR reads. HC = Haplotype Coverage, ER = Error Rate (mismatches + indels + ‘N’s). MC = Misassembled contigs proportion. NGA50 is labeled with ‘-’ if the uniquely aligned blocks cover less than half of the reference length. *If contigs are full-length, this number represents the estimated number of haplotypes or strains in the virus mixture. The total sequencing coverage in this table is 20000×.

|  | #Contigs* | HC (%) | N50 | NGA50 | ER (%) | MC(%) |
| --- | --- | --- | --- | --- | --- | --- |
| *5-strain HIV mixture* |  |  |  |  |  |  |
| Strainline | 5 | 99.9 | 9697 | 9697 | 0.002 | 0.0 |
| Canu | 5 | 84.5 | 8227 | 8170 | 0.409 | 20.0 |
| wtdbg2 | 1 | 15.5 | 7419 | - | 1.820 | 0.0 |
| *6-strain poliovirus mixture* |  |  |  |  |  |  |
| Strainline | 6 | 99.9 | 7449 | 7444 | 0.074 | 0.0 |
| Canu | 6 | 62.7 | 7040 | 6399 | 0.538 | 0.0 |
| wtdbg2 | 1 | 14.7 | 6575 | - | 0.244 | 0.0 |
| *10-strain HCV mixture* |  |  |  |  |  |  |
| Strainline | 10 | 99.9 | 9294 | 9292 | 0.056 | 0.0 |
| Canu | 11 | 76.9 | 7703 | 7174 | 0.351 | 0.0 |
| wtdbg2 | 2 | 13.6 | 7698 | - | 5.077 | 0.0 |
| *15-strain ZIKV mixture* |  |  |  |  |  |  |
| Strainline | 15 | 99.6 | 10238 | 10238 | 0.021 | 0.0 |
| Canu | 13 | 55.7 | 10233 | 7129 | 0.189 | 0.0 |
| wtdbg2 | 2 | 10.7 | 8773 | - | 1.693 | 0.0 |
| *5-strain SARS-CoV-2 mixture* |  |  |  |  |  |  |
| Strainline | 7 | 98.6 | 29017 | 29009 | 0.047 | 0.0 |
| Canu | 16 | 85.9 | 12419 | 25137 | 0.078 | 0.0 |
| wtdbg2 | 1 | 20.6 | 29158 | - | 0.360 | 0.0 |

#### Simulated ONT data

Table 2 displays the benchmarking results for Oxford Nanopore reads assembly. Again, Strainline yields near-perfect results on all five datasets. All true haplotypes are reconstructed (HC≈100%, recall=100%) with extremely low error rate (0.023% ~ 0.081%), and without misassemblies. Moreover, it achieves the exact number of true strains on three datasets (HIV, Poliovirus and HCV, precision=100%), and overestimates the number of haplotypes by one on the 15-strain ZIKV and the 5-strain SARS-CoV-2 mixtures (precision is 100% and 83.3%, respectively).

**Table 2.** Benchmarking results for simulated Oxford Nanopore reads. HC = Haplotype Coverage, ER = Error Rate (mismatches + indels + ‘N’s). MC = Misassembled contigs proportion. NGA50 is labeled with ‘-’ if the uniquely aligned blocks cover less than half of the reference length. The total sequencing coverage in this table is 20000×.

|  | #Contigs | HC (%) | N50 | NGA50 | ER (%) | MC(%) |
| --- | --- | --- | --- | --- | --- | --- |
| *5-strain HIV mixture* |  |  |  |  |  |  |
| Strainline | 5 | 99.9 | 9702 | 9702 | 0.081 | 0.0 |
| Canu | 8 | 58.1 | 9088 | 9086 | 1.066 | 0.0 |
| wtdbg2 | 1 | 18.9 | 9046 | - | 1.327 | 0.0 |
| *6-strain poliovirus mixture* |  |  |  |  |  |  |
| Strainline | 6 | 100.0 | 7454 | 7453 | 0.051 | 0.0 |
| Canu | 1 | 16.6 | 7448 | - | 0.781 | 0.0 |
| wtdbg2 | - | - | - | - | - | - |
| *10-strain HCV mixture* |  |  |  |  |  |  |
| Strainline | 10 | 99.9 | 9294 | 9294 | 0.023 | 0.0 |
| Canu | 1 | 10.0 | 9280 | - | 2.511 | 0.0 |
| wtdbg2 | 2 | 18.4 | 8567 | - | 1.336 | 0.0 |
| *15-strain ZIKV mixture* |  |  |  |  |  |  |
| Strainline | 16 | 98.3 | 10244 | 10244 | 0.026 | 0.0 |
| Canu | 1 | 6.6 | 10251 | - | 0.459 | 0.0 |
| wtdbg2 | 3 | 17.1 | 9490 | - | 0.564 | 0.0 |
| *5-strain SARS-CoV-2 mixture* |  |  |  |  |  |  |
| Strainline | 6 | 99.9 | 29299 | 29071 | 0.041 | 0.0 |
| Canu | 10 | 66.3 | 18003 | 9492 | 0.062 | 0.0 |
| wtdbg2 | 1 | 19.0 | 26767 | - | 0.586 | 0.0 |

In comparison, Canu obtains 16.6% 10.0% and 6.6% haplotype coverage on Poliovirus, HCV, ZIKV datasets, respectively, which indicates that Canu does not operate in a strain-specific manner. On HIV and SARS-CoV-2, Canu achieves better haplotype coverage, while still missing a considerable proportion of strains (HC = 58% and 66%, respectively). While Canu has 100% recall on SARS-CoV-2, only 0% to 20% recall are achieved on the other four datasets (see Figure S2). Canu also shows relatively high error rates in the HIV and HCV assemblies (1.0% and 2.5%; 13 and 109 times higher than Strainline, respectively). For Poliovirus and ZIKV datasets, Canu displays about 15 times higher error rates in comparison with Strainline.

Wtdbg2 only yields 17% to 19% haplotype coverage on four datasets (except Poliovirus) at relatively high error rates (e.g. 1.3% on HIV and HCV). Wtdbg2 failed to run on the Poliovirus dataset, so no results are shown.

#### Real data

See Table 3 for results on the 5-strain PVY mixture. Also here, Strainline reconstructs the great majority of strain-specific sequences (HC=97.9%, recall=60%). Importantly, recall is 100% if edit distance is set to 3. Similarly, Strainline overestimates the number of haplotypes by two, but achieves perfect precision (100%) when operating at edit distance 3.

**Table 3.** Benchmarking results for real Oxford Nanopore reads. HC = Haplotype Coverage, ER = Error Rate (mismatches + indels + ‘N’s). MC = Misassembled contigs proportion. NGA50 is labeled with ‘-’ if the uniquely aligned blocks cover less than half of the reference length. * Sometimes NGA50 still reports a value (5456 bp) even if HC< 50% because contigs have overlaps (See https://github.com/ablab/quast/discussions/174 for the detailed explanation). Note that the metrics for the SARS-CoV-2 real sample in this table are not necessarily correct but for reference only, because the ground truth is unknown and we only used the sequence of Wuhan-Hu-1 (NC 045512) as the truth for comparison.

|  | #Contigs | HC (%) | N50 | NGA50 | ER (%) | MC(%) |
| --- | --- | --- | --- | --- | --- | --- |
| *5-strain PVY mixture* |  |  |  |  |  |  |
| Strainline | 7 | 97.9 | 9538 | 9548 | 0.956 | 0.0 |
| Canu | 3 | 39.9 | 9665 | 5456* | 0.105 | 0.0 |
| wtdbg2 | 2 | 26.0 | 7632 | - | 4.931 | 0.0 |
| *SARS-CoV-2 (SRP250446)* |  |  |  |  |  |  |
| Strainline | 1 | 99.9 | 29565 | 29565 | 0.832 | 0.0 |
| Canu | - | - | - | - | - | - |
| wtdbg2 | 1 | 65.8 | 19405 | 19396 | 1.542 | 0.0 |

Canu only reconstructs the most abundant two strains at full coverage and high accuracy (error rate for these two strains is 0.07% and 0.09%). However, Canu misses to cover any of the other three strains (reflected by HC=40% and recall=40%). Strainline, on the other hand, reconstructs the most abundant two strains at their full length and the exact same low error rate as Canu. In addition, unlike Canu, Strainline assembles also the other less abundant three strains at full coverage and sufficiently low error rate (0.18% to 2.7%). Wtdbg2 only generates one single near full-length haplotype and another one short contig (HC=26%) at an error rate of 4.9%.

As for the SARS-CoV-2 real sample (SRP250446), we use the genome sequence of Wuhan-Hu-1 (NC 045512) as the reference for comparison since the ground truth is unknown. Strainline yields one single full-length haplotype (HC=99.9%) at an error rate clearly below the sequencing error rate (0.8%). Wtdbg2 only obtains one fragmented contig (HC=65.8%, N50=19405) with about two times higher error rate (1.5%). Canu was unable to finish after running for more than ten days on a 48-core computing machine, clearly exceeding acceptable computational resources, so we stopped the job. This explains why no results are shown.

### Haplotype abundance estimation

We also evaluated the accuracy of estimated haplotype abundances on both simulated and real data, see Table 4. Note that Canu and Wtdbg2 do not provide abundance estimation (because of their general setup as consensus assemblers, so no comparison with other methods can be provided.

**Table 4.** Absolute and relative errors of estimated haplotype abundances by Strainline on different virus mixtures. For each dataset, we present the average error over all assembled strains. Note that for the *SARS-CoV-2 (SRP250446)* real sample, the abundance estimation error is not shown because the ground truth is unknown. AFE: absolute frequency error. RFE: relative frequency error.

| Datasets | AFE (%) | RFE (%) |
| --- | --- | --- |
| *Simulated PacBio CLR* |  |  |
| 5-strain HIV | 0.04 | 0.18 |
| 6-strain poliovirus | 2.17 | 48.35 |
| 10-strain HCV | 0.06 | 0.80 |
| 15-strain ZIKV | 0.18 | 4.93 |
| 5-strain SARS-CoV-2 | 3.40 | 17.49 |
| *Simulated ONT* |  |  |
| 5-strain HIV | 0.27 | 1.48 |
| 6-strain poliovirus | 2.30 | 51.18 |
| 10-strain HCV | 0.28 | 3.05 |
| 15-strain ZIKV | 0.28 | 6.10 |
| 5-strain SARS-CoV-2 | 4.14 | 20.60 |
| *Real ONT* |  |  |
| 5-strain PVY | 4.48 | 23.07 |

Strainline estimates the frequencies for the reconstructed haplotypes at operable accuracy, with absolute frequency error (AFE) of 0.04%/0.27% (PacBio / ONT) on the simulated HIV data, 0.06%/0.28% on the HCV data and 0.18%/0.28% on the ZIKV data, respectively. One observes that the relative frequency errors (RFE) follow a similar pattern. On simulated Poliovirus, SARS-CoV-2 and real Poliovirus, Strainline estimates the abundances at moderate accuracy (AFE is between 2.17% and 4.48%). A likely explanation for increased error rates on Poliovirus data is the fact that strain abundances vary exponentially (exponentially increasing from 1.6% to 50.8%). The less accurate estimates on Poliovirus refer mainly to low frequent strains, which naturally causes high relative frequency errors (48%, 51% on PacBio and ONT data). A possible reason for increased abundance estimates error on the simulated SARS-CoV-2 and the real PVY datasets is the overestimation of the number of haplotypes in the mixed samples. These findings suggest that accurate haplotype reconstruction goes hand in hand with accurate haplotype abundance estimation. The results are also likely to reflect the current limits in that respect, because coverage fluctuations and nevertheless shorter reads impose certain constraints on estimating haplotype abundance.

### Effect of sequencing coverage

To investigate the effect of sequencing coverage on viral quasispecies assembly, we chose a 5-strain HIV mixture, as one of the most challenging datasets suggested in Baaijens et al. (2017, 2019). We simulated PacBio CLR reads with different overall sequencing coverage. Assembly results are shown in Table 5. We observe that Strainline successfully reconstructs all true haplotypes at nearly perfect coverage (HC > 99%) and great accuracy sequencing coverage exceeds 2000×. Strainline outperforms all other methods substantially in this respect. On decreasing sequencing depth (i.e. 1000 to 500×), Strainline does not reconstruct all haplotypes at perfect coverage, but stays at HC approaching 80%. Still, Strainline clearly outperforms the other tools in terms of haplotype coverage (HC ranges from 14% to 64%), N50 and NGA50. Also, Strainline has similar accuracy like Canu.

**Table 5.** Benchmarking results for 5-strain HIV mixture (PacBio CLR reads) with varying sequencing coverage. HC = Haplotype Coverage, ER = Error Rate (mismatches + indels + ‘N’s). MC = Misassembled contigs proportion.

|  | #Contigs | HC (%) | N50 | NGA50 | ER (%) | MC(%) |
| --- | --- | --- | --- | --- | --- | --- |
| *500×* |  |  |  |  |  |  |
| Strainline | 7 | 78.4 | 9517 | 9545 | 1.603 | 0.0 |
| Canu | 4 | 32.9 | 6766 | 5281 | 1.447 | 0.0 |
| wtdbg2 | 1 | 18.0 | 8687 | - | 2.295 | 0.0 |
| *1000×* |  |  |  |  |  |  |
| Strainline | 5 | 79.2 | 9557 | 9554 | 1.069 | 0.0 |
| Canu | 5 | 63.6 | 7207 | 7139 | 1.087 | 0.0 |
| wtdbg2 | 1 | 13.5 | 6516 | - | 1.520 | 0.0 |
| *2000×* |  |  |  |  |  |  |
| Strainline | 6 | 99.3 | 9644 | 9614 | 0.675 | 0.0 |
| Canu | 5 | 67.4 | 8193 | 7544 | 0.416 | 20.0 |
| wtdbg2 | 1 | 16.0 | 7741 | - | 1.103 | 0.0 |
| *5000×* |  |  |  |  |  |  |
| Strainline | 5 | 99.7 | 9690 | 9686 | 0.311 | 0.0 |
| Canu | 4 | 69.3 | 8264 | 8001 | 0.537 | 25.0 |
| wtdbg2 | 1 | 16.5 | 7950 | - | 2.115 | 0.0 |
| *10000×* |  |  |  |  |  |  |
| Strainline | 5 | 99.8 | 9696 | 9691 | 0.184 | 0.0 |
| Canu | 4 | 71.7 | 8844 | 8524 | 0.446 | 25.0 |
| wtdbg2 | 1 | 15.6 | 7528 | - | 2.777 | 0.0 |
| *20000×* |  |  |  |  |  |  |
| Strainline | 5 | 99.9 | 9697 | 9697 | 0.002 | 0.0 |
| Canu | 5 | 84.5 | 8227 | 8170 | 0.409 | 20.0 |
| wtdbg2 | 1 | 15.5 | 7419 | - | 1.820 | 0.0 |

As the sequencing coverage increases, both Strainline and Canu achieve better assemblies in terms of haplotype coverage and error rate (which of course is no surprise). wtdbg2 however generates one single consensus sequence of high error content (1.1% to 2.8%) in all cases. Interestingly, when raising sequencing depth beyond 2000×, the accuracy of Canu does not improve, with the error rate staying between 0.4% and 0.5%. Canu also introduces one misassembly at greater depth. However, as for our method, increasing sequencing coverage always reduce the error rates of the assemblies. Specifically, specifically, declines by nearly two orders of magnitude when depth is raised from 10000× to 20000×.

### Runtime and memory usage evaluation

We performed all benchmarking analyses on a x86 64 GNU/Linux machine with 48 CPUs. The runtime and peak memory usage evaluations for different methods are reported in Supplementary Tables S1 and S2. Undoubtedly, wtdbg2 is the fastest tool thanks to the efficiency of fuzzy de Bruijn graphs, which only takes a few seconds and 0.1~0.4 GB memory on all datasets. For simulated PacBio CLR reads assembly, Strainline is 1.4~16 times faster and requires less or similar peak memory in comparison to Canu (Table S1). For simulated ONT reads assembly, Strainline is 1.8 to 8 times slower and requires 5 to 9 times more memory than Canu on HIV, Poliovirus and HCV data (Table S2). However, on ZIKV and SARS-CoV-2, Canu is 15 to 38 times slower and requires about 6 times more memory in comparison with Strainline (Table S2). By design, the most expensive steps of Strainline are threaded, and there are two steps that consume time, namely ‘Correction’ and ‘Consensus’, see Figure 1. Overall, Strainline requires 12 to 108 CPU hours and 15 to 44 GB main memory on the datasets of 20000× coverage. This indicates that our method is very well applicable in all real world scenarios of interest.

## Discussion

We have presented Strainline, an approach that reconstructs haplotype-specific genomes from mixed viral samples using noisy long-read sequencing data. To the best of our knowledge, Strainline is the first such approach that is presented in the literature.

Although the length of long reads is a major advantage in the assembly of genomes, the greatly elevated error rates pose substantial challenges when seeking to distinguish between little diverse genomes. The large amount of sequencing errors that affect the reads easily exceed the amount of genetic variants that are characteristic of the different genomes. Because co-occurring true mutations can mean a decisive handle in the differential analysis, it is usually advantageous to make use of the reads at their full length. However, addressing this particular challenge by computing all-vs-all overlaps of reads may demand excessive runtimes.

To address these major challenges, we proceed by drawing from de Bruijn graph based and overlap graph based techniques in a way that aims to optimally combine the virtues of the two paradigms. We first employ a local de Bruijn graph based strategy by way of an initial error correction step. Remarkably, this strategy was originally been designed and presented for correcting long reads sampled from prokaryotic and eukaryotic genomes, without that the strategy established the state of the art on such longer genomes. Here, we realized that it performs just perfectly fine when considering short genomes. The stragegy successfully suppresses sequencing errors to a degree that enables us to carry out the second step.

The second step clusters the pre-corrected long reads into groups of reads that are supposed to collect reads from identical haplotypes. To avoid excessive overlap computations, we iteratively select seed reads, as longest reads that do not overlap any of the previously selected seed reads. If reads sufficiently overlap a seed read, we put them into the cluster of the respective seed read. This way, we determine clusters based on seed-vs-all overlap computations. Because the number of seed reads is smaller by orders of magnitude in comparison with the number of reads overall, we reduce the runtime by orders of magnitude in comparison with performing all-vs-all overlap computations. Subsequently, we generate a haplotype-specific consensus (haplotig) for each cluster of reads, which further eliminates errors and preserves haplotype-specific variation. Upon having computed this consensus, we iteratively extend the haplotigs by evaluating their overlaps—note that the number of haplotigs is much smaller than the number of reads, such that all-vs-all haplotig overlap computations are easily feasible. In a last step, we discard haplotigs (haplotypes) of too low divergence or abundance. The final output is a reliable set of (often full-length) haplotypes together with their abundances. In this, the goal of *de novo viral quasispecies assembly from noisy long reads has been achieved*.

Benchmarking experiments on both simulated and real data, reflecting various mixed viral samples referring to various relevant settings, such as different viruses, different numbers of strains, haplotype abundances and sequencing platforms (PacBio/ONT), have shown that our approach accurately reconstructs the haplotype-specific sequences. Thereby, the output contigs tend to cover all haplotypes at their full length.

As a result, Strainline proves superior in comparison with all methods that are currently available when trying to assemble virus genomes from long read data. The superiority of Strainline becomes documented in terms of various well-known and -approved assembly evaluation metrics: Strainline’s contigs cover substantially more haplotypes, are longer (N50, NGA50) and are more accurate in terms of error and misassembly rates. Of course, Strainline’s superiority did not come as a particular surprise, because Strainline has been the first approach to address the challenge upfront, while the competitors do not decidedly address it. While Canu at least is able to recover a certain fraction of haplotypes, wtdbg2 always outputs a single consensus sequence. Because Canu and wtdbg2 had been the only approaches that one could use, Strainline arguably establishes a major step up in the haplotype-specific assembly of viral genomes when using noisy third-generation sequencing reads.

There were some further clear hints that our approach made sense. First, we demonstrated that Strainline was able to exploit increasing sequencing coverage to its advantage; note that deeply sequenced datasets are common when analyzing virus genomes. At the same time, Strainline required the least amount of reads for establishing sufficiently accurate haplotypes in comparison with other methods. This indicates that Strainline caters to a greater range of experimental settings of practical interest.

In addition, Strainline is the only approach available that does not only assemble the viral genomes, but also estimates the abundances of the haplotypes that make part of the mix of viral strains. Results have demonstrated that Strainline’s abundance estimates are sufficiently accurate if the abundances do not refer to strains of very low relative frequencies. Note however that low-frequency strains pose particular challenges not only in this respect, because the relative lack of coverage for such strains raises the level of uncertainty one has to deal with.

Note finally that in comparison with short-read viral quasispecies assemblers (such as most prominently Baaijens et al. (2019, 2020)) that accurately operate in a haplotype-specific way, Strainline is the only approach that reconstructs the haplotypes in all datasets at their full length. This does not only point out that long reads indeed do mean a major advantage over short reads, but also means that Strainline is able to leverage the advantages of long reads successfully.

Nonetheless, improvements are conceivable. Sometimes, Strainline tends to overestimate the number of haplotypes, which as a consequence inevitably hampers the estimation of the abundances of the strains. One possible reason for haplotype overestimation is that haplotype identity in read overlaps is based on overlap length and sequence identity alone, which may be too simplistic. Likely, more sophisticated criteria will be able to mend this issue, which we consider valuable future work. In addition, the computational efficiency of the approach has further room for improvement. For example, the computation of consensus sequence from read clusters can be implemented in a more efficient way.

## Conclusions

This paper presents Strainline, an approach to full-length viral haplotype reconstruction from noisy long-read sequencing data. Strainline operates in a *de novo* fashion, that is, Strainline does not make use of a reference at any time. We make use of local De Bruijn graph assembly to sufficiently correct sequencing errors in raw reads, such that it is possible to extend contigs iteratively at a haplotype-specific level, in order to eventually yield full-length strain-specific haplotypes. These properties render Strainline unique in the spectrum of currently available assemblers: it is the only approach that can reconstruct the strain-specific haplotypes in mixed viral samples using long reads, and accurately estimate their abundances.

We remain with saying that databases (e.g. GISAID) are currently filling up with SARS-CoV-2 TGS sequencing read samples, drawn from infected people. So far, one has been blind with respect to counting the number of strains that commonly affect their hosts in an unbiased way. The biased view on the amount of strains that have infected people initially, or have formed during the course of the infection is a decisive hindrance when assessing the evolutionary development of SARS-CoV-2. Seeing the full spectrum of strains, without having to make use of existing, potentially already obsolete reference genomes, has the potential to yield major insight into the course of the pandemic. Now, we can finally have a closer look. Our approach is implemented in an easy-to-use open-source tool https://github.com/xiaoluo91/Strainline.

## Methods

### Correcting sequencing errors

For initial correction of errors in long reads, we adopt a local De Bruijn graph assembly based strategy. While de Bruijn graph based data structures tend to have difficulties when dealing with TGS data because of the high error rates, it is shown in Tischler and Myers (2017) that it can work effectively when applied to small segments of the long reads.

Here, we realized that the corresponding strategy is particularly powerful when applied to virus TGS data. In our experiments, we observed that the local de Bruijn graph based strategy has substantial advantages on virus data in comparison to the results presented in the seminal work Tischler and Myers (2017), which exclusively focused on TGS data from prokaryotic and eukaryotic genomes of length at least a few Mbp. We further realized that we could use Daccord, the corresponding tool Tischler and Myers (2017), by straightforward modification of a few parameters.

The main steps of the workflow that address error correction are shown in Figure 2. In a first step (see “Target read & overlapping reads”), read overlaps of raw reads are computed using Daligner V2.0 Myers (2014), which uses canonical k-mers to identify significant local alignments between reads. Upon having selected a target read, we subsequently (see “Read alignment pile”) form a read alignment pile, consisting of the target read and all reads that share significant overlap with it. Then (see “Windows”), we divide the pile into small windows, which serves the application of the local de Bruijn graph based strategy; note that the windows have to be sufficiently small (here: 40 bp) such that the strategy works satisfyingly.

Accordingly, we construct de Bruijn graphs for all windows of size 40 bp (see “DBGs for all windows”). Importantly, windows may share a small overlapping interval, as one can see in “Window consensus”: windows and their overlapping intervals can be interpreted as nodes and edges of another graph. The respective graph of windows and overlapping intervals can then be traversed, where scores can be assigned to paths through that graph. The paths of windows through that graph that are optimal in terms of the scores are determined, and are further evaluated with respect to differences with the target read; the concatenation of sequences of the graph that has highest score and least differences in comparison with the target read is taken as the consensus sequence, reflects the true sequence that underlies the target read. One can then correct the errors in the target read accordingly.

### Read cluster generation

Subsequently, we compute clusters of (error corrected) overlapping reads. This addresses to wipe out further errors, and, as the major point, to form groups of reads all of which stem from the same (local) haplotype. For pseudo code supporting the generation of read clusters, see Algorithm 1.

For this, first, we sort the error corrected reads by length in decreasing order, considering that longer reads tend to have more overlaps. Processing reads in the corresponding order therefore results in larger read clusters. This increases the length of the resulting haplotigs and hence improves the assembly overall. In each iteration, we choose the longest read having remained unprocessed as the seed read and compute seed-vs-all overlaps on corrected reads using Minimap2 Li (2018), whose seed-chain-align procedure is known to perform pairwise alignments extremely fast. Bad overlaps are filtered out according to reasonable, additional criteria. For example, overlaps that are too short, that do not exceed a minimum level of sequence identity, that reflect self-overlaps, duplicates or internal matches are removed.

In this, we follow Algorithm 5 in Li (2016). Enforced by a strict threshold, the remaining overlapping reads are expected to stem from the haplotype of the seed read. The corresponding cluster is determined as the set of reads that overlap the seed read (according to the criteria listed above).

Subsequently, all of the reads of the cluster are discarded from the sorted list of reads, and the next iteration (referring to line 5 in Algorithm 1) is executed. The procedure stops when the number of iterations (hence clusters) reaches the upper limit *k* where *k* is user defined (default 100), or all reads have been processed (corresponding to *R* = in Algorithm 1).

Notably, we only compute seed-vs-all overlaps, and not all-vs-all overlaps (as per, for example, a straightforward, naive approach), and limit the number of clusters, which decisively speeds up the procedure. This procedure is supported by the fact that, on the one hand, it is common that the coverage for virus sequencing data ultra-deep, and, on the other hand, that the majority of sequencing errors in the raw long reads have already been corrected in the error correction step. An all-vs-all overlap computation would have to deal with massive amounts of overlapping reads, which is unnecessary.

**Algorithm 1.**
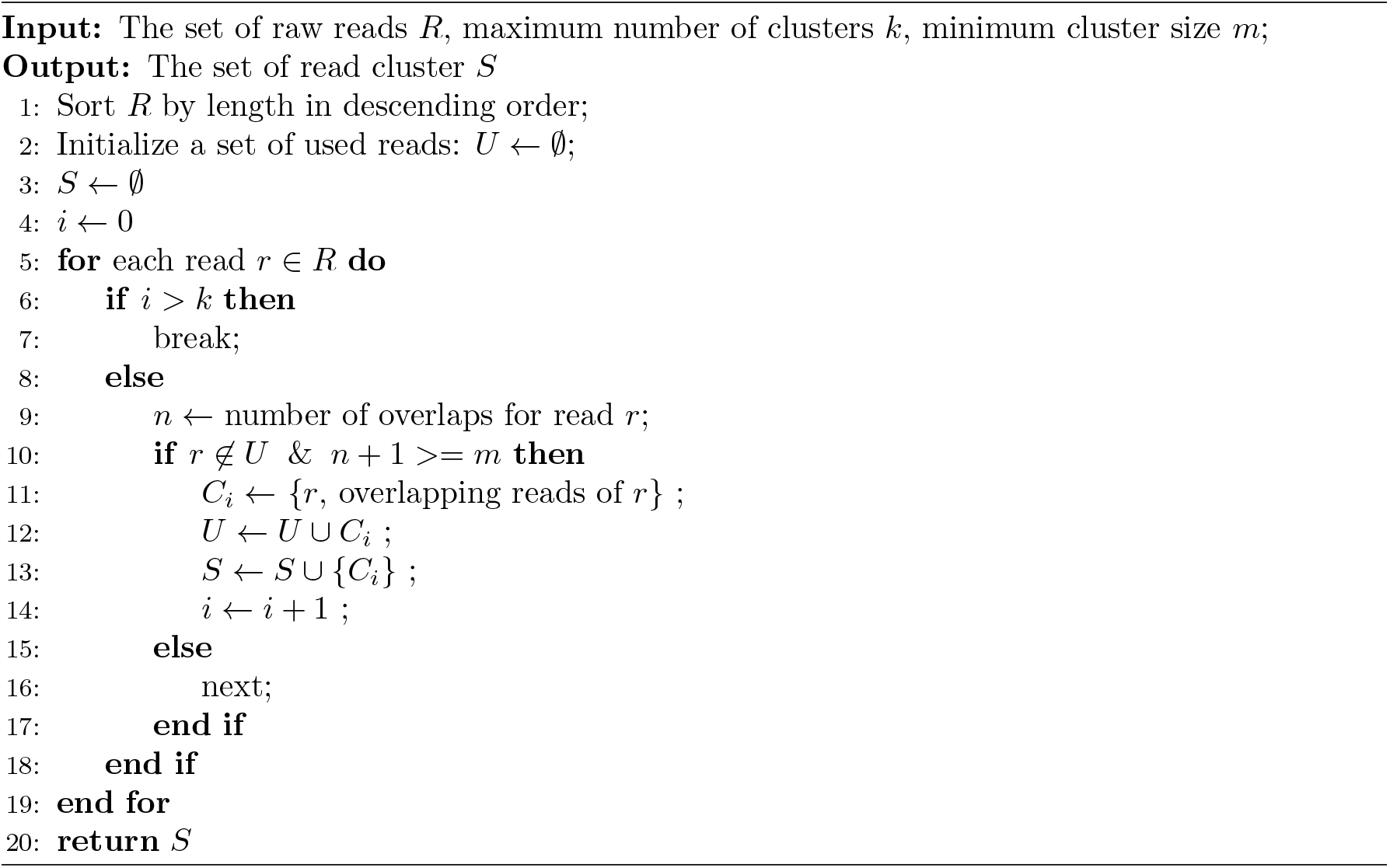
Compute read clusters

### Generating a consensus sequence for each read cluster

Although reads were initially corrected, they may still contain errors. The major possible reasons are near-identical genomic regions that are shared across haplotypes.

For final polishing of reads, and removal of also more stubbornly resisting errors, we first compute a partial order alignment (POA) algorithm Lee et al. (2002) for each cluster. Subsequently, we generate the consensus sequence for the POA of each cluster (which is a straightforward, generic procedure). Adapting reads to this consensus completes the process of error correction.

For computing POA’s of clusters, we make use of the fast SIMD version, as implemented in Spoa Vaser et al. (2017), and built into our approach. Note that this step can generate a longer and more accurate haplotype-specific sequence for each cluster (which reflects the cluster-specific haplotig).

### Iterative extension of haplotigs

Haplotigs generated from the previous step do not necessarily reflect full-length haplotypes. This explains why it makes sense to try to extend them further.

For extending haplotigs, one considers haplotigs as reads, an re-runs “Read cluster generation” and “Generating a consensus sequence for each read cluster” another time, inserting the haplotigs of the first iteration as reads. The procedure is iteratively repeated until haplotigs cannot be elongated further. In that, our experiments demonstrate that two iterations usually suffice for virus sequencing data sets.

### Haplotype filtration

Ideally, iteratively extending haplotigs eventually results in correct, full-length haplotypes. However, in practice, it is possible that some haplotypes have very low pairwise divergence or very low relative abundance, both of which indicates that the corresponding haplotypes reflect artifacts. It is therefore reasonable to filter out such artificial haplotypes, because they either reflect redundant or spurious sequences. For the identification of low divergence and low relative abundance haplotypes, we make use of two procedures for computing haplotype divergence on the one hand, and haplotype relative abundance on the other hand.

#### Haplotype divergence calculation

We propose two metrics for haplotype divergence measurement, namely local divergence (LD) and global divergence (GD). Given two haplotypes *Hi, Hj*, let *l* be the length of their overlap, and *m* be the number of identically matching positions in the overlap (so *m* ≤ *l* by definition of *l, m*). Let further *ni, nj* be the lengths of the non-overlapping parts of *Hi, Hj* relative to their overlap.

LD is defined by the formula *LD*(*H_i_, H_j_*) = 1 − *m/l* and GD is defined by *GD*(*H_i_, H_j_*) = 1 − *m/*(*l* + *n_i_* + *n_j_*). In other words, LD agrees with BLAST-like alignment identity, when only considering the overlapping regions. GD, on the other hand, considers the entire sequence context that neighbors and includes the overlap of *H_i_* and *H_j_*.

Note that two haplotypes having low local divergence but large global divergence (because of a long overhang) are more likely to stem from two different strains than haplotypes having small LD and GD.

Let further *maxLD* and *maxOH* represent the user-defined maximum local divergence and the maximum overhang length (5bp in our cases), respectively. Note that if *H_i_* being contained in *H_j_* implies length of *H_i_* being less than the length of *H_j_*, as well as *LD*(*H_i_, H_j_ < maxLD* and overhang length at most *maxOH*. In this case, *H_i_* is discarded from the downstream analysis. For determining the overlap information of two haplotypes, Minimap2 is used. The ultimate output is a set of non-redundant haplotypes.

#### Calculating relative abundance of haplotypes

Calculating haplotype relative abundance is straightforward when the length of the haplotypes approaches the size of the (strain-specific) genomes, and when original reads are nearly free of errors.

For the calculation, one aligns the reads to the haplotypes, which given the situation, can usually be done in a unique, non-ambiguous way. The result of the alignments is stored in a BAM file. We then simply adopt the `jgi_summarize_ bam_contig_depths` program from MetaBAT 2 *Kang et al.* (2019) to calculate the depth of haplotypes based on the BAM file. The relative abundance of *H_i_* is equal to the average depth of *H_i_* divided by the overall average depth of all haplotypes. Haplotypes with very low relative abundance are filtered out, and one recomputes the abundance for the remaining haplotypes upon removal of the spurious, low abundance haplotypes.

The final output consists of a set of full-length haplotypes along with their corresponding relative frequencies, as desired.

### Data simulation

To evaluate performance of Strainline, we generated several simulated datasets for both PacBio CLR and ONT reads. For simulating reads, we made use of PBSIM Ono et al. (2013) model-based simulation, which reflects a sound way to generate PacBio CLR reads of N50 length 2.4kbp and average sequencing error rate 10%. In addition, we also downloaded real Oxford Nanopore reads (GridION) of a SARS-CoV-2 sample from the SRA database: SRP250446 and then used NanoSim V2.6.0 *Yang et al.* (2017) as an approved simulator to train an ONT read profile based on this real ONT dataset. Accordingly, we generated simulated ONT reads, at an N50 of 2.5kbp in terms of length and average sequencing error rate of 10%. The genomes used for each dataset are listed in the Availability of data and materials.

## Acknowledgements

We would thank Jasmijn Baaijens for the insightful discussions.

## Availability of data and materials

All data (including raw sequencing reads, reference genomes and assemblies) and code for reproducing the results in the paper are deposited in Code Ocean with the capsule DOI https://doi.org/10.24433/CO.3155281.v1. The source code of Strainline is GPL-3.0 licensed, and publicly available at https://github.com/xiaoluo91/Strainline.

## Ethics approval and consent to participate

Not applicable.

## Consent for publication

Not applicable.

## Authors’ contributions

XL and AS developed the method. XL, XK, and AS wrote the manuscript. XL and XK conducted the data analysis. XL implemented the software. All authors read and approved the final version of the manuscript.

## Footnotes

1 Receiving the error message “No disjointigs were assembled”; upon contact, the authors responded that metaFlye does not support the assembly of very short sequences, such as viruses.

## References

Baaijens, J.A. et al (2017). De novo assembly of viral quasispecies using overlap graphs. Genome research, 27(5), 835–848.

Baaijens, J.A. et al (2019). Full-length de novo viral quasispecies assembly through variation graph construction. Bioinformatics, 35(24), 5086–5094.

Baaijens, J.A., Stougie, L. and Schönhuth, A. (2020). Strain-aware assembly of genomes from mixed samples using flow variation graphs. In International Conference on Research in Computational Molecular Biology, pages 221–222. Springer.

Beerenwinkel, N. et al (2005). Computational methods for the design of effective therapies against drug resistant hiv strains. Bioinformatics, 21(21), 3943–3950.

Chin, C.S. et al (2016). Phased diploid genome assembly with single-molecule real-time sequencing. Nature methods, 13(12), 1050–1054.

Della Bartola, M., Byrne, S. and Mullins, E. (2020). Characterization of potato virus y isolates and assessment of nanopore sequencing to detect and genotype potato viruses. Viruses, 12(4), 478.

Domingo, E. et al (1996). Basic concepts in rna virus evolution. The FASEB Journal, 10(8), 859–864.

Domingo, E., Sheldon, J. and Perales, C. (2012). Viral quasispecies evolution. Microbiology and Molecular Biology Reviews, 76(2), 159–216.

Douek, D.C., Kwong, P.D. and Nabel, G.J. (2006). The rational design of an aids vaccine. Cell, 124(4), 677–681.

Freire, B. et al (2021). Inference of viral quasispecies with a paired de bruijn graph. Bioinformatics, 37(4), 473–481.

Giallonardo, F.D. et al (2014). Full-length haplotype reconstruction to infer the structure of heterogeneous virus populations. Nucleic acids research, 42(14), e115–e115.

Holland, J.J.d., De La Torre, J. and Steinhauer, D. (1992). Rna virus populations as quasispecies. Genetic diversity of RNA viruses, pages 1–20.

Kang, D.D. et al (2019). Metabat 2: an adaptive binning algorithm for robust and efficient genome reconstruction from metagenome assemblies. PeerJ, 7, e7359.

Knyazev, S. et al (2021). Epidemiological data analysis of viral quasispecies in the next-generation sequencing era. Briefings in bioinformatics, 22(1), 96–108.

Kolmogorov, M. et al (2019). Assembly of long, error-prone reads using repeat graphs. Nature biotechnology, 37(5), 540–546.

Koren, S. et al (2017). Canu: scalable and accurate long-read assembly via adaptive k-mer weighting and repeat separation. Genome research, 27(5), 722–736.

Lee, C., Grasso, C. and Sharlow, M.F. (2002). Multiple sequence alignment using partial order graphs. Bioinformatics, 18(3), 452–464.

Li, H. (2016). Minimap and miniasm: fast mapping and de novo assembly for noisy long sequences. Bioinformatics, 32(14), 2103–2110.

Li, H. (2018). Minimap2: pairwise alignment for nucleotide sequences. Bioinformatics, 34(18), 3094–3100.

Logsdon, G.A., Vollger, M.R. and Eichler, E.E. (2020). Long-read human genome sequencing and its applications. Nature Reviews Genetics, 21(10), 597–614.

Loman, N.J. et al (2013). A culture-independent sequence-based metagenomics approach to the investigation of an outbreak of shiga-toxigenic escherichia coli o104: H4. Jama, 309(14), 1502–1510.

Mikheenko, A. et al (2018). Versatile genome assembly evaluation with quast-lg. Bioinformatics, 34(13), i142–i150.

Myers, G. (2014). Efficient local alignment discovery amongst noisy long reads. In International Workshop on Algorithms in Bioinformatics, pages 52–67. Springer.

Ono, Y., Asai, K. and Hamada, M. (2013). Pbsim: Pacbio reads simulator—toward accurate genome assembly. Bioinformatics, 29(1), 119–121.

Prabhakaran, S. et al (2013). Hiv haplotype inference using a propagating dirichlet process mixture model. IEEE/ACM transactions on computational biology and bioinformatics, 11(1), 182–191.

Ruan, J. and Li, H. (2020). Fast and accurate long-read assembly with wtdbg2. Nature methods, 17(2), 155–158.

Shafin, K. et al (2020). Nanopore sequencing and the shasta toolkit enable efficient de novo assembly of eleven human genomes. Nature Biotechnology, pages 1–10.

Somerville, V. et al (2019). Long-read based de novo assembly of low-complexity metagenome samples results in finished genomes and reveals insights into strain diversity and an active phage system. BMC microbiology, 19(1), 1–18.

Tischler, G. and Myers, E.W. (2017). Non hybrid long read consensus using local de bruijn graph assembly. bioRxiv, page 106252.

Töpfer, A. et al (2014). Viral quasispecies assembly via maximal clique enumeration. PLoS Comput Biol, 10(3), e1003515.

Vaser, R. et al (2017). Fast and accurate de novo genome assembly from long uncorrected reads. Genome research, 27(5), 737–746.

Vicedomini, R. et al (2021). Automated strain separation in low-complexity metagenomes using long reads. bioRxiv.

Yang, C. et al (2017). Nanosim: nanopore sequence read simulator based on statistical characterization. GigaScience, 6(4), gix010.

Zagordi, O. et al (2011). Shorah: estimating the genetic diversity of a mixed sample from next-generation sequencing data. BMC bioinformatics, 12(1), 1–5.

