## Supplementary Material for "Strainline: full-length de novo viral haplotype reconstruction from noisy long reads"

### Other benchmarking tools

Canu version: snapshot (), installed on Nov4, 2019 via conda
Wtdbg2 version: 0.0 (19830203)

| | CPU time (h) | peak memory usage (GB) |
|---|---|---|
| *5-strain HIV mixture* | | |
| Strainline | 18.9 | 20.6 |
| Canu | 313.5 | 49.5 |
| Wtdgb2 | 0.002 | 0.4 |
| *6-strain poliovirus mixture* | | |
| Strainline | 39.8 | 14.9 |
| Canu | 239.4 | 52.8 |
| Wtdgb2 | 0.001 | 0.1 |
| *10-strain HCV mixture* | | |
| Strainline | 52.0 | 20.0 |
| Canu | 311.8 | 29.4 |
| Wtdgb2 | 0.002 | 0.1 |
| *15-strain ZIKV mixture* | | |
| Strainline | 99.7 | 22.3 |
| Canu | 143.8 | 16.3 |
| Wtdgb2 | 0.001 | 0.1 |
| *5-strain SARS-CoV-2 mixture* | | |
| Strainline | 107.7 | 44.1 |
| Canu | 667.3 | 87.5 |
| Wtdgb2 | 0.006 | 0.1 |

**Table S1.** Runtime and memory usage for simulated PacBio CLR reads assembly. The total sequencing coverage in this table is 20000×.

| | CPU time (h) | peak memory usage (GB) |
|---|---|---|
| *5-strain HIV mixture* | | |
| Strainline | 11.9 | 19.4 |
| Canu | 3.2 | 2.2 |
| Wtdgb2 | 0.001 | 0.4 |
| *6-strain poliovirus mixture* | | |
| Strainline | 24.1 | 19.2 |
| Canu | 3.2 | 2.3 |
| Wtdgb2 | 0.001 | 0.1 |
| *10-strain HCV mixture* | | |
| Strainline | 17.6 | 17.9 |
| Canu | 10.2 | 3.7 |
| Wtdgb2 | 0.001 | 0.1 |
| *15-strain ZIKV mixture* | | |
| Strainline | 77.0 | 16.5 |
| Canu | 1181.4 | 95.2 |
| Wtdgb2 | 0.001 | 0.1 |
| *5-strain SARS-CoV-2 mixture* | | |
| Strainline | 29.6 | 18.9 |
| Canu | 1134.0 | 127.4 |
| Wtdgb2 | 0.005 | 0.2 |

**Table S2.** Runtime and memory usage for simulated ONT reads assembly. The total sequencing coverage in this table is 20000×.

**(a)**

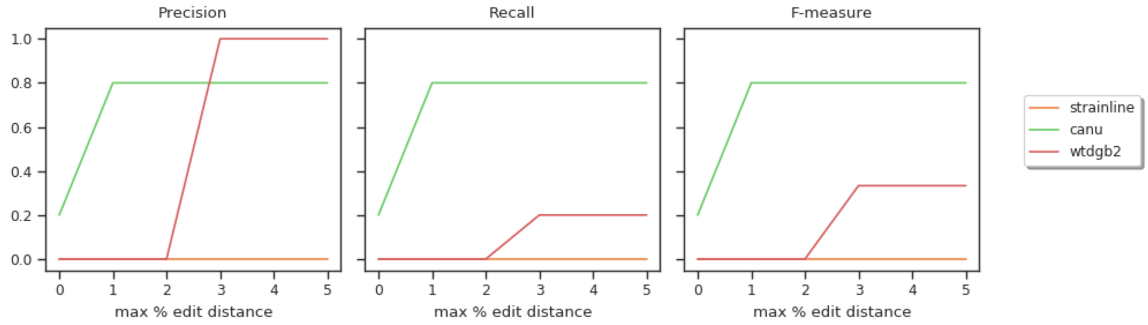
Precision
Recall
F-measure
max % edit distance
max % edit distance
max % edit distance
strainline
canu
wtdgb2

**(b)**

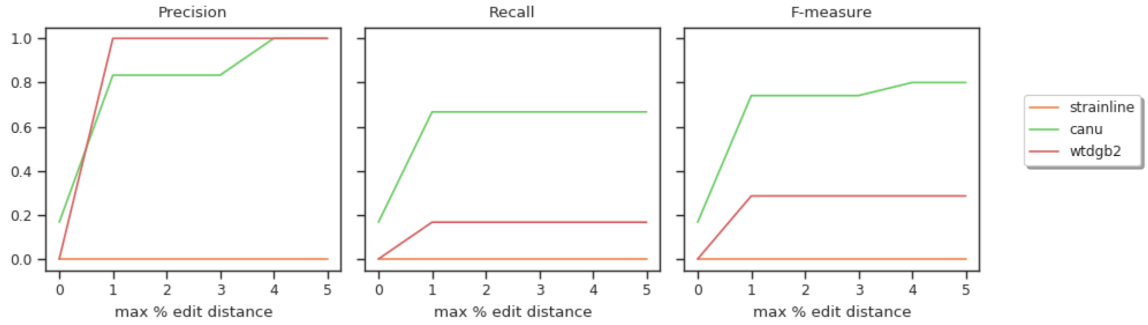
Precision
Recall
F-measure
max % edit distance
max % edit distance
max % edit distance
strainline
canu
wtdgb2

**(c)**

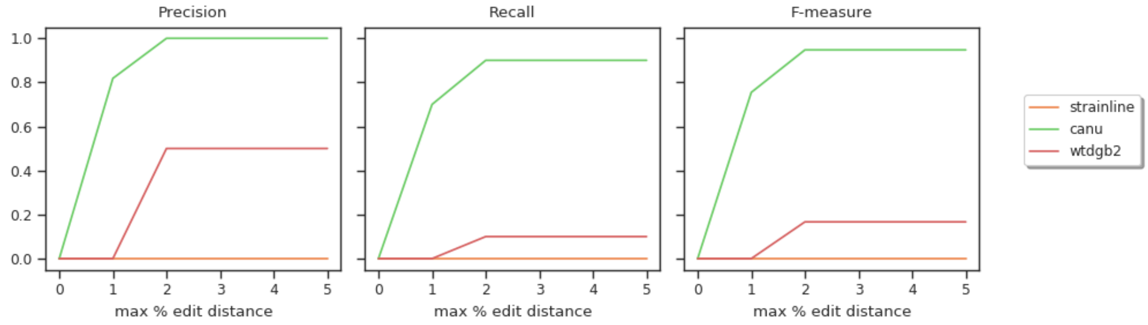
Precision
Recall
F-measure
max % edit distance
max % edit distance
max % edit distance
strainline
canu
wtdgb2

**(d)**

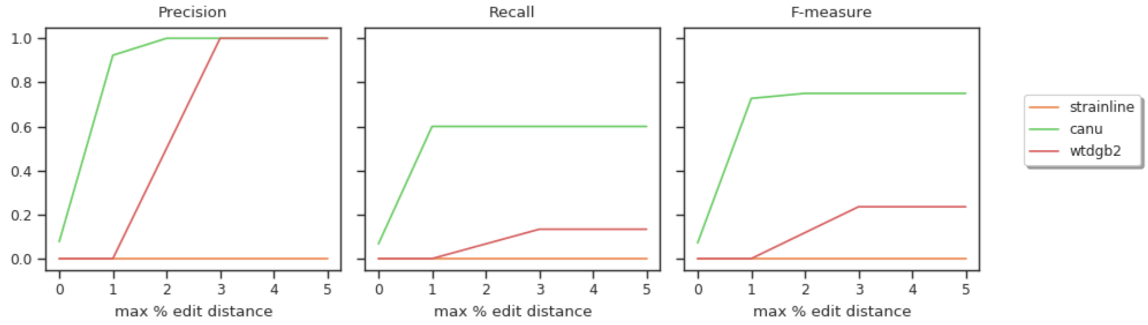
Precision
Recall
F-measure
max % edit distance
max % edit distance
max % edit distance
strainline
canu
wtdgb2

**(e)**

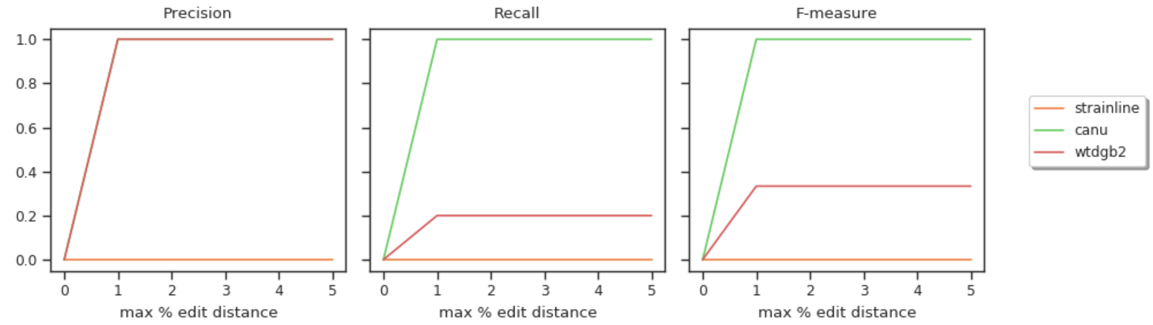


**Figure S1.** Precision, recall and F-measure for different viral mixtures (simulated PacBio CLR reads). **a,b,c,d,e** represent 5-strain HIV, 6-strain Poliovirus, 10-strain HCV, 15-strain ZIKV, 5-strain SARS-CoV-2 mixtures, respectively. The x-axis denotes the various thresholds (max edit distance) which are used to determine if an assembled contig is assigned to a true strain.

**(a)**

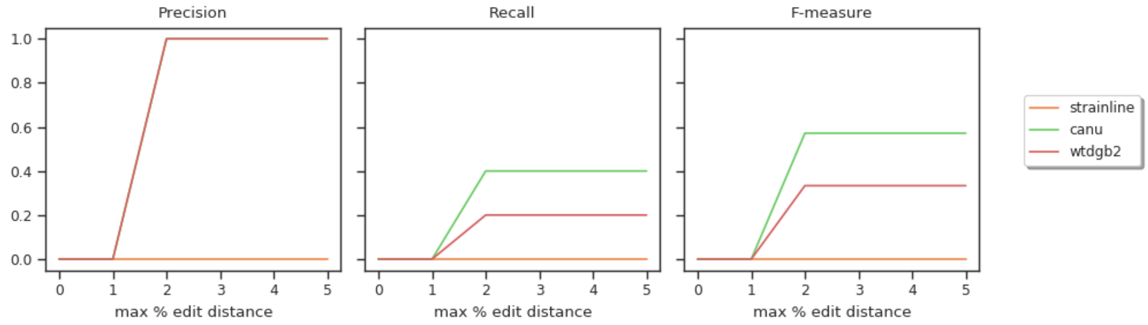
Precision
Recall
F-measure
1.0
0.8
0.6
0.4
0.2
0.0
0
1
2
3
4
5
max % edit distance
strainline
canu
wtdgb2

**(b)**

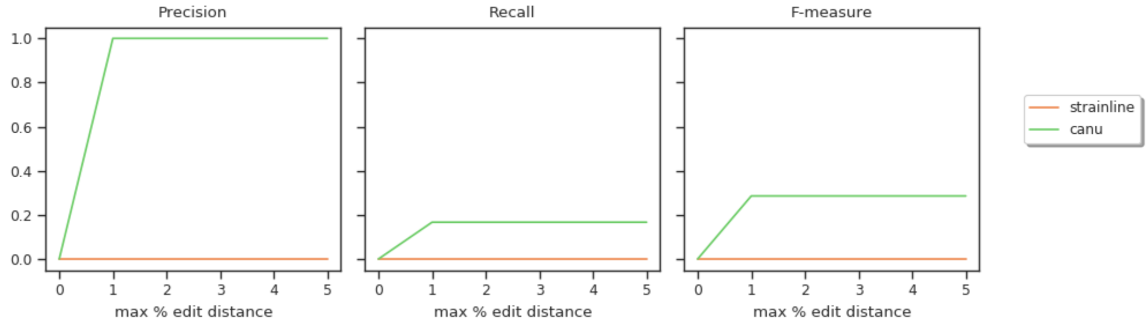
Precision
Recall
F-measure
1.0
0.8
0.6
0.4
0.2
0.0
0
1
2
3
4
5
max % edit distance
strainline
canu

**(c)**

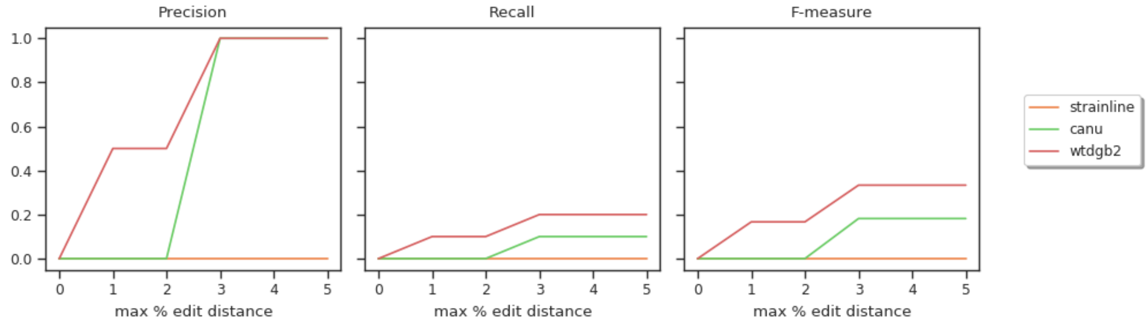
Precision
Recall
F-measure
1.0
0.8
0.6
0.4
0.2
0.0
0
1
2
3
4
5
max % edit distance
strainline
canu
wtdgb2

**(d)**

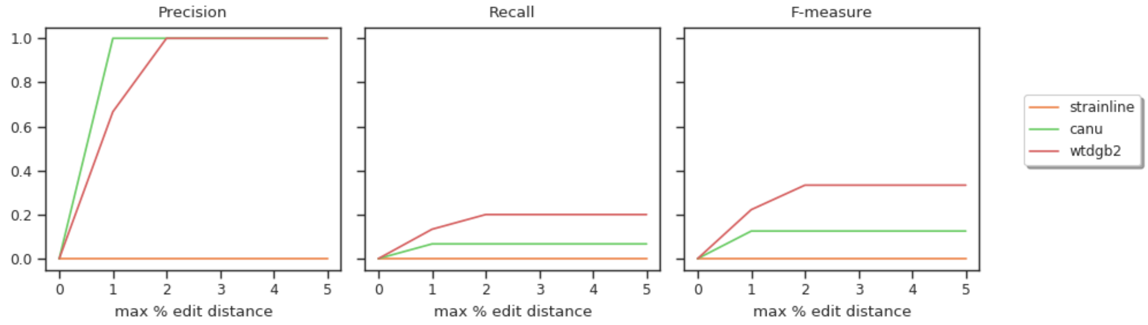
Precision
Recall
F-measure
1.0
0.8
0.6
0.4
0.2
0.0
0
1
2
3
4
5
max % edit distance
strainline
canu
wtdgb2

(e)

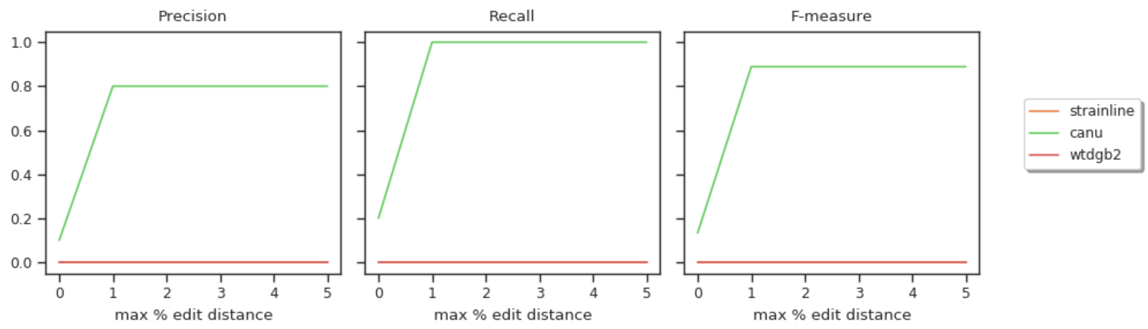


**Figure S2.** Precision, recall and F-measure for different viral mixtures (simulated Oxford Nanopore reads). **a,b,c,d,e** represent 5-strain HIV, 6-strain Poliovirus, 10-strain HCV, 15-strain ZIKV, 5-strain SARS-CoV-2 mixtures, respectively. The x-axis denotes the various thresholds (max edit distance) which are used to determine if an assembled contig is assigned to a true strain.

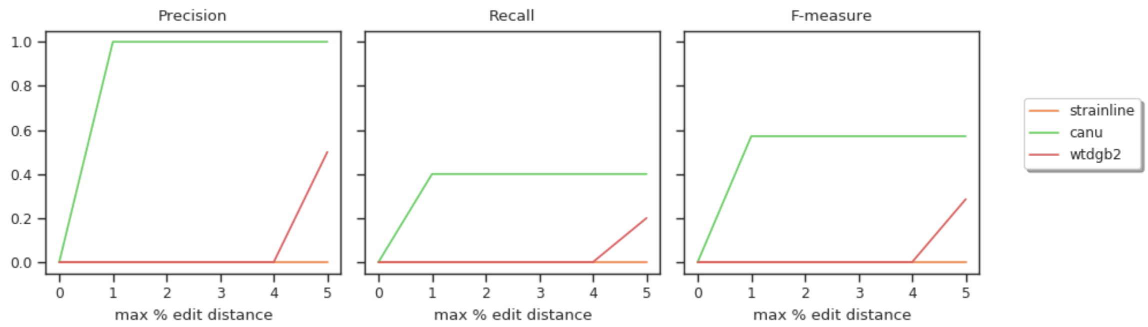


**Figure S3.** Precision, recall and F-measure for 5-strain PVY mixture (real data). The x-axis denotes the various thresholds (max edit distance) which are used to determine if an assembled contig is assigned to a true strain.
